# Tie2 controls the dynamics of tight junction assembly

**DOI:** 10.64898/2026.11.23.768735

**Authors:** YT Zhao, DD Ehnes, AR Hayes, P Panfil, A Phal, SY Zhu, C Hoffman, Xinru Wang, A Morishita, D Baker, J Mathieu, H Ruohola-Baker

## Abstract

The Angiopoietin-Tie2 pathway is a master regulator of vascular homeostasis and stability. This study demonstrates Tie2 regulates tight junction formation at both transcriptional and molecular level. Tie2 receptor clusters trigger the formation of two classes of Tie2 complexes. The Tie2-α5β1 integrin complex activates pAKT/FOXO1 signaling that promotes the gene expression of tight junctions. Independent of α5β1, Tie2 drives the direct recruitment and assembly of tight junctions—including ZO1, claudin-5, and occludin—to endothelial cell-cell junctions. In iPSC-derived endothelial cell differentiation, Tie2 activation upregulates tight junction ZO1 formation at the cell-cell junction, thereby promoting endothelial maturation. In a diabetic blood vessel organoid model, targeting the Tie2 pathway successfully rescues key pathological phenotypes by reducing collagen IV expression and restoring tight junction integrity. Our findings establish that Tie2 activation is a crucial driver of tight junction assembly in endothelial cells, offering a potent therapeutic strategy for treating diabetic vasculopathy and other conditions characterized by vascular instability.

**Significance statement:** While the angiopoietin-Tie2 pathway has long been recognized as one of the key regulators of vascular development during embryogenesis and postnatal vascular remodeling, the molecular basis of how Tie2 orchestrates this process remains unclear. By utilizing designed protein constructs, we dissected this process and demonstrated evidence showing that Tie2 clustering orchestrates vascular stability through two specialized signaling complexes. We show that Tie2 activation by high-valency ligands serves as a critical driver of tight junction assembly, directly recruiting essential proteins like ZO1, claudin-5, and occludin to endothelial junctions independent of α5β1 integrin, which, to our knowledge has not been described before. While Tie2-α5β1 complex regulates pAKT/FOXO1 and promotes the gene expression of tight junctions. Our data illustrated a novel mechanism of how Tie2 directly regulates vascular stability and a new therapeutic target for vascular diseases.

## INTRODUCTION

Vascular stability is maintained through the coordinated activity of VEGF, FGF, and Tie2 signaling pathways, together with integrins and tight junction proteins. The angiopoietin-Tie2 pathway regulates blood vessel stability, remodeling, and permeability(1–4). Angiopoietin 1 (Ang1) promotes pAKT activation and blood vessel stabilization, while Angiopoietin 2 (Ang2) inhibits pAKT and induces leaky vasculatures despite sharing nearly identical protein structures in receptor-binding domains (5–8). Previously, we demonstrated using designed protein scaffolds that Ang1 and Ang2 activities fall into two broad phenotypic classes distinguished by the number of presented Ang1 F-domains that describe the mechanism of Tie2 signaling output(9–11). We found that ligands presenting six or more copies of F-domains behave like Ang 1, activating pAKT, while trimers behave like Ang2, inhibiting Tie2-induced pAKT.

α5β1 is a heterodimeric integrin family receptor that regulates cell adhesion and migration via focal adhesion complexes(12–14). α5β1 is upregulated in newly formed blood vessels(14), and genetic ablation of α5 integrin in mice is embryonic lethal due to severe vascular defects and leakage(15, 16). Activation of α5β1 using fibronectin has been shown to promote α5β1-Tie2 interaction and enhance Ang1-dependent pERK1/2 activation(17, 18). We have previously shown that 2-dimensional protein sheets presenting high copies of F-domain (Fd-sheet) promote Tie2 co-localization with α5β1 integrin, and this association is crucial for pAKT activation(9). In contrast, others also show that inhibition of α5β1 integrin promoted Ang2-mediated Tie2 activation (19). The mode of α5β1-Tie2 interaction and signaling output correlates with downstream functional cellular outcome.

Tight junction proteins are critical regulators of vascular stability(20–24). Following ischemic brain injury in mice, temporally coordinated increases in angiopoietin-1 (Ang1), integrin α5, claudin-5, and ZO-1 expression at sites of vascular disruption suggest that Ang1–Tie2 signaling and α5β1 integrin may contribute to blood–brain barrier (BBB) repair (25–27). However, other studies suggest that α5β1 can compromise vascular stability (28, 29) with α5 inhibition enhancing BBB integrity and increasing claudin-5 expression ((28, 29)). Disruption of α5β1 has also been associated with increased localization of VE-cadherin (CD144) and ZO-1 at cell–cell junctions (19, 30). Together, these findings highlight the need to further clarify how α5β1, Tie2, and tight junction proteins interact to control vascular stability.

In this study, we investigated the molecular mechanisms governing vascular stability, focusing on the interplay between Tie2, α5β1 integrin, and the tight junction proteins ZO-1 and claudin-5. Using *de novo* designed protein scaffolds and binders, we identified two distinct mechanisms by which Tie2 regulates vascular stability. High-valency ligands induce Tie2 clustering and promote junctional recruitment of ZO-1, claudin-5, and occludin independently of α5β1 integrin, whereas the Tie2–α5β1 complex regulates AKT–FOXO1 signaling and tight junction gene expression. These findings reveal complementary roles for Tie2 in junction assembly and transcriptional regulation, suggesting new therapeutic strategies for vascular diseases.

## RESULTS

### Tie2 recruits tight junctions in an integrin-independent manner

We have previously shown that clustering Tie2 receptors with high F-domain valency ligands induced the colocalization with α5 integrin subunit(9). In this study, we investigated the interactions between Tie2, α5β1 integrin, and tight junction proteins to examine vascular dynamics by using a micron-scale protein array tagged with GFP and conjugated with Ang1 F-domain (Fd-sheet(31)) to cluster Tie2 receptors (Fig. 1A). To investigate Tie2 interactions with integrin and tight junction molecules, serum-starved HUVECs were treated with the Fd-sheets for 30 minutes before PFA fixation and immunofluorescence staining for microscopy analysis. Upon clustering Tie2, we found that α5 and active β1 integrins, along with the downstream target, pCAS, colocalized with Tie2 under the Fd-sheets, while inactive β1 is not observed (Fig. 1B-D & Fig. S1A-C). Our data suggest clustering of Tie2 receptors using Ang 1 F-domains can recruit and induce α5β1 activation, illustrated by the presence of active β1 and pCas colocalizing with Tie2. We then analyzed the presence of tight junction markers upon clustering Tie2 using the Fd-sheets. Our data show that Tie2 co-localizes with ZO1, CLDN5, and OCLN (Fig. 1E-G and S1D).

**Figure 1:**
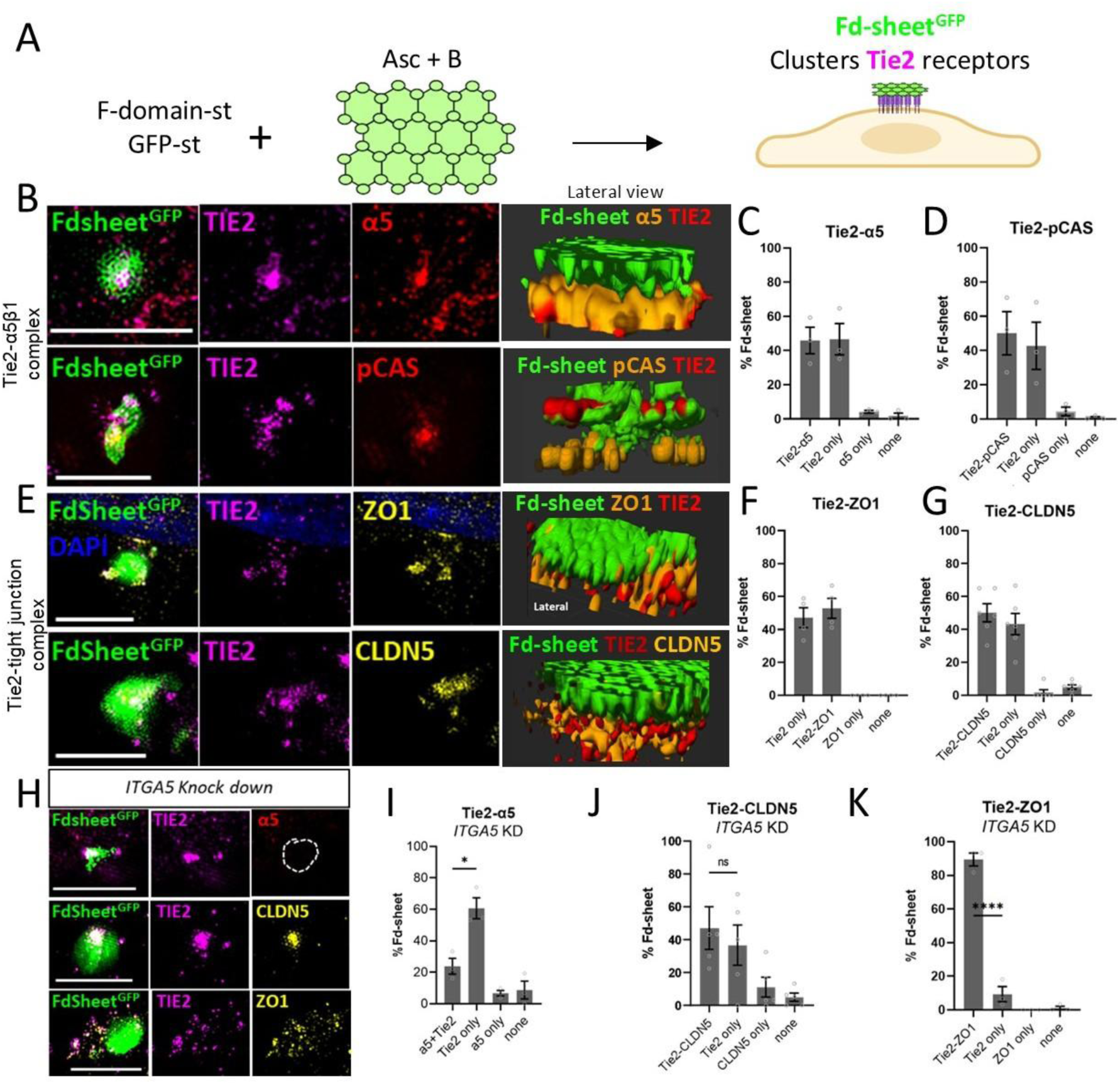
Two classes of Tie2 complexes. **A)** F-domain fused with SpyTag (Fd-st) and GFP-st were conjugated to two-dimensional protein sheets to make Fd-sheets for clustering Tie2 receptors on serum-starved HUVECs for 30 minutes before fixation for immunofluorescence staining. Figure S1C shows a histogram describing the proportion of F-domain sheets shows Tie2 colocalization in HUVECs. **B & E)** Super-resolution OMX images of immunofluorescence staining (left 3 panels) and imaris 3D reconstruction (right gray panels) of Tie2 colocalizing with α5, pCas, ZO1, or claudin-5 (CLDN5), forming two classes of Tie2 complexes: Tie2-α5/pCAS and Tie2-ZO1/CLDN5. Figure S1E shows a zoomed-out image of the Tie2-pCAS cluster. **C-D & F-G)** Quantifications of the different Tie2 complexes under the Fd-domain sheets. **H)** Super-resolution OMX images of immunofluorescence staining of HUVECs pre-treated with siRNA against *ITGA5* before serum starvation, and then Fd-sheet administration. **I-K)** Quantifications showing the changes in the composition of the Tie2 complexes upon *ITGA5* knockdown. As negative controls, we showed that Tie2 clusters do not colocalize with unrelated receptors, notch, GRP161, and α3 integrin as well as negative Tie2 regulator, VE-PTP, is also absent in Tie2 clusters (Fig S1P-R). A Two-tailed t-test was performed to evaluate the level of statistical significance; *p*-value < 0.05 is annotated with *, *p*-value < 0.0001 is annotated with ****, and ns means not significant. Each data point is a biological repeat for all experiments and at least 30 Fd-sheets were counted in each biological replicate. All scale bars are 5 µm.

α5β1 and Tie2 were previously suggested to play a role in tight junction formation for vascular repair(27); thus, we investigated whether the Tie2-α5β1 complex regulates the recruitment of tight junction molecules. Using 5-color confocal microscopy, we stained cells for CLDN5, Tie2, and active β1 (Fig. S1F). Interestingly, our quantifications revealed two classes of Tie2 complexes: Tie2-active β1 or Tie2-CLDN5, with minimal overlap of the two complexes (Fig. S1G). To further evaluate whether α5β1 regulates the formation of the Tie2-tight junction complex, we knocked down *ITGA5* using siRNA before treating the cells with Fd-sheets (Fig. S1H). Our results show that inhibiting α5β1 via α5 knock down did not affect the recruitment of ZO1 and CLDN5 to Tie2 clusters (Fig. 1H-K). Our finding is consistent with studies showing that α5β1 inhibition did not affect endothelial tight junction formation (15, 19, 29, 30). Collectively, we demonstrated that Tie2 activation recruits tight junction proteins in an integrin-independent manner.

Further investigation of α5 knock down revealed no changes in the portion of Fd-sheet binding to cells since Fd-sheet can still bind to Tie2 (Fig S1J-J). Whereas knocking down Tie2 reduced Fd-sheet binding to cells, and the proportion of Fd-sheet binding to cells via α5 increased compared to wild type (Fig. S1I-K). Using the fluorescence polarization assay, the fluorescence signal generated by FITC-tagged cRGD binding to α5β1 integrin decreases when H8F is added, indicating a competitive binding mode in which FITC-cRGD is competed away due to H8F binding to α5β1 integrin (Fig. S1L). Taken together, our data are consistent with previous studies showing that the F-domain can also bind α5β1 integrin in the absence of Tie2(32, 33).

Adherens junctions such as VE-cadherin (CD144) also play critical roles in vascular stability and permeability(20, 34). Adherens junction formation at the paracellular space is a key prerequisite for the formation of tight junctions. A recent study shows inactive β1 integrin colocalizes with VE-cadherin, serving as a scaffold that stabilizes VE-cadherin at the cell-cell junction(35). Interestingly, we observed that inhibiting α5β1 using siRNA significantly reduced the level of Tie2-VE-cadherin colocalization compared to wild-type control, suggesting that the Tie2-α5β1 complex may be important for initiating cell-cell adhesion prior to tight junction formation (Fig. S1M-O).

In summary, upon clustering Tie2 receptors, we observed two classes of Tie2 complexes: Tie2-α5β1 and Tie2-tight junctions (CLDN/OCLN/ZO1). Integrin is not needed for Tie2-mediated tight junction recruitment but may play a role in tight junction stability. We then investigated the downstream functional outcome of these complexes.

### The Tie2-α5β1 complex regulates pAKT/FOXO1 for tight junction gene expression

We previously demonstrated that α5β1 is essential for Tie2/pAKT activation, as siRNA-mediated knockdown of α5 robustly inhibits this signaling cascade(9). To further probe this mechanism, we employed α5β1mb (36)—a monomeric, synthetic de novo mini-binder targeting the integrin ligand-binding interface—to competitively block fibronectin or F-domain binding. Serum-starved HUVECs were treated with the Tie2 super-agonists H8F(9) to activate Tie2/pAKT signaling, and downstream FOXO1 translocation out if the nucleus to promote cell survival(37, 38) (Fig. 2A-B & Fig. S2). However, when α5 expression was knocked down via siRNA, the capacity of H8F to induce FOXO1 nuclear exclusion was significantly impaired (Fig. 2C-D). Similarly, co-treatment with H8F and α5β1mb resulted in FOXO1 nuclear retention (39). In contrast, inhibiting other integrins using de novo mini-binders targeting αvβ6 and αvβ8 (14, 40) in competition with Tie2 agonist, Icos1(9) did not affect pAKT activity or FOXO1 translocation (Fig. S2B-E). Together, these findings establish that α5β1 is specifically required to drive the Tie2/pAKT/FOXO1 signaling axis.

**Figure 2:**
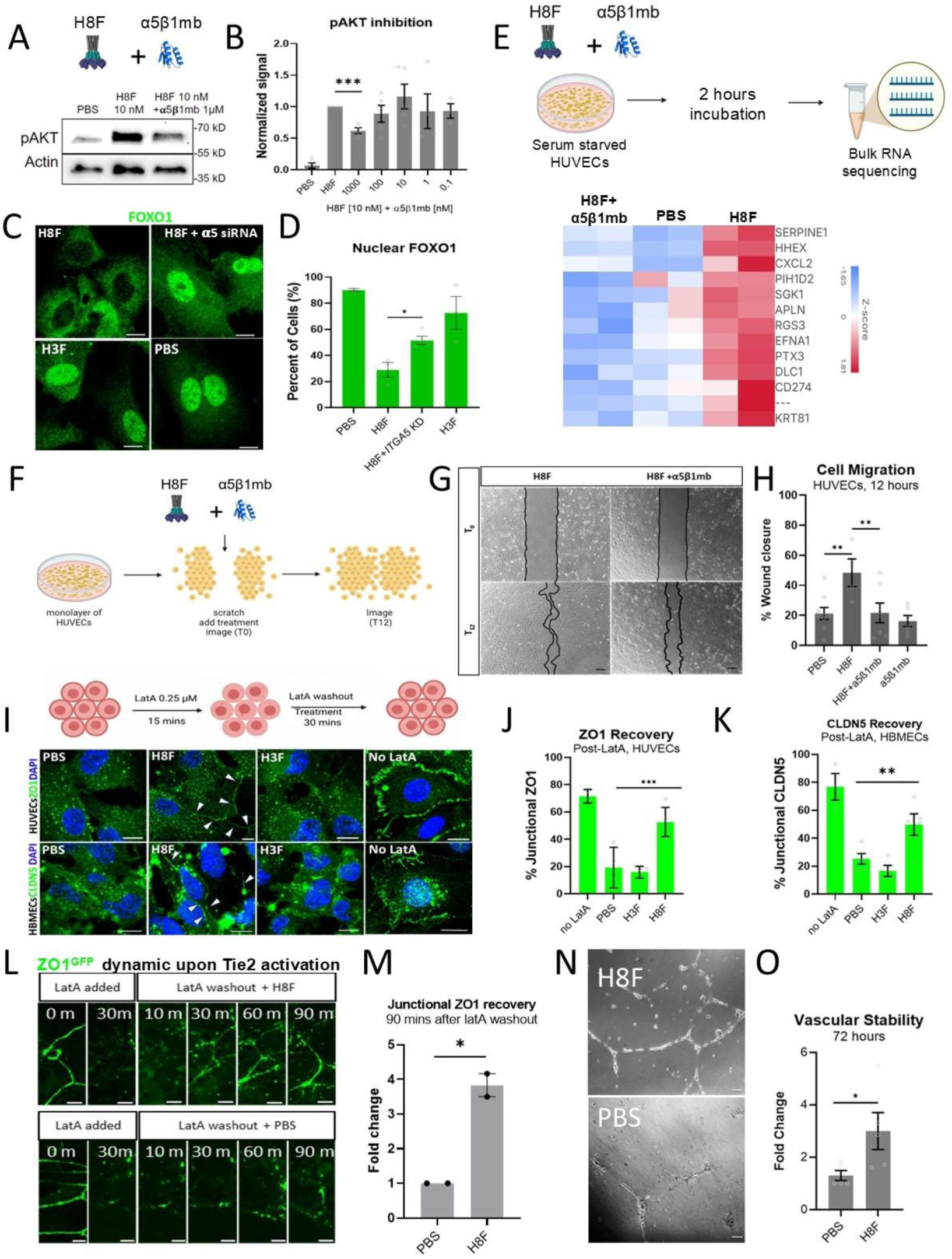
Tie2 forms different complexes to regulate cell migration or tight junctional assembly. **A)** Competition experiments of serum-starved HUVECs were treated with 10 nM of H8F with or without 1µM of α5β1mb, and protein lysates were collected for western blot staining to analyze pAKT activity. **B)** Quantification of pAKT level from western gels. pAKT signal was normalized to actin and then normalized to H8F samples. **C)** Confocal images of FOXO1 immunofluorescence staining of serum-starved HUVECs treated with 50 nM of H8F +/- 1µM of α5β1mb or PBS or HUVECs pre-treated with siRNA before starvation and H8F administration, scale bars are 10 µm. **D)** Quantification of FOXO1 images, at least 100 cells were counted in each biological replicate. **E)** Top: schematic of bulk RNA sequencing experiment. Serum-starved HUVECs were treated with H8F 10nM, H8F 10nM + α5β1mb 1µM, or PBS for two hours before RNA lysate was collected for RNA sequencing. Bottom: bulk RNA sequencing data were analyzed using Partek DESeq2 (Fig. S3A), a portion of the heatmap of differentially expressed genes in the green box is shown. **F)** Schematic of scratch assay in which a wound was made in the middle of a monolayer of HUVECs before adding H8F 10nM +/- α5β1mb 500nM for 12 hours. **G)** Representative brightfield images of scratched area at 0 and 12-hour timepoints, scale bars are 200 µm. **H)** Images of cells at 0- and 12-hour timepoints were quantified to evaluate the level of cell migration using ImageJ. Each data point is a biological replicate. **I)** Top: schematic of Lat-A induced tight junction disassembly and recovery assay. bottom: confocal images of ZO1 or claudin-5 (CLDN5) immunofluorescence staining of HUVECs or HBMECs, respectively, after LatA washout and recovery was supplemented with 100 nM of H8F, H8T, H3T, or PBS, scale bars are 10 µm. **J-K)** Quantification showing the percentage of cells stained with a junctional ZO1 or claudin-5 lining, at least 100 cells were counted in each biological replicate. **L)** Live-cell images of iPSC-derived day 14 endothelial cells with ZO1 genetically tagged with GFP. iENDOs were treated with 0.5 µM of Lat-A for 15 minutes and then washed once with PBS before adding H8F and imaged up to 1.5 hours, scale bars are 5 µm. **M)** Quantification of junctional ZO1 at 90 minutes timepoint images using Gemini 3.1 Pro. **N)** Brightfield images of HUVECs at the 72-hour timepoint showing the formation and stability of the tubule network, bars are 100 µm. **O)** Quantifications of the tubule network at 72 hours. Images were quantified using the ImageJ angiogenesis analyzer plugin. The number of tubes, meshes, and nodes was averaged to calculate vascular stability and then normalized to PBS control. One-way ANOVA with Bonferroni post hoc test was used to analyze level of significance, *p*-value < 0.05, 0.01, 0.0001 are annotated *, **, ****, respectively. Each data point is a biological repeat for all experiments.

The pAKT/FOXO1 pathway regulates the transcriptional expression of genes that regulate vascular stability(37, 40), thus we investigated whether inhibiting integrin in the Tie2-α5β1 complex using α5β1mb would downregulate genes related to vascular stability. Serum starved HUVECs were treated with H8F, H8F + α5β1mb, or PBS for 2 hours before cell lysates were collected for bulk RNA sequencing. RNA sequencing data were analyzed for DESeq2 and filtered for significantly downregulated genes in H8F+α5β1mb samples compared to H8F samples (FDR <0.05 and p-value <0.05) (Fig. S3A). The significantly downregulated genes were analyzed for pathway enrichment. As expected, genes related to the pAKT/FOXO1 pathway were enriched in downregulated genes when comparing H8F+α5β1mb to H8F (Fig. S3B), indicating that H8F activates the Tie2-α5β1 complex for vascular stability but inhibiting α5β1 downregulates vascular stability. Specific genes related to vascular stability such as DLC1(41, 42), HHEX(43–46), APLN(47), and SGK1(48, 49), the primary drivers that promote vascular stability, integrity, or mature vessel homeostasis were upregulated after H8F treatment, but significantly downregulated when α5β1 was inhibited (H8F+ α5β1mb) (Fig. 2E). Notability, HHEX is a key transcriptional regulator that preserves endothelial quiescence by downregulating VEGFA, VEGFR1, and VEGFR2 (43, 46, 50) to prevent excessive VEGF signaling from triggering vascular remodeling. These data show that the Tie2-α5β1 complex regulates vascular stability at the transcriptional level via the pAKT/FOXO signaling axis.

### Tie2-α5β1 complex regulates cell migration

In Figure 1, we showed that the Tie2-α5β1 recruits pCAS (Fig. 1B-E), a key component in the focal adhesion complex that regulates cell migration(51, 52). To validate the role of the Tie2-α5β1 complex in endothelial cell migration, we used α5β1mb with H8F in the scratch assay to analyze Tie2-induced cell migration(9, 36, 53). Briefly, a scratch was made in the monolayer of HUVECs, which were then treated with Tie2 agonists (H8F or Icos1) with or without α5β1mb for 12 hours to evaluate the change in wound area (Fig. 2F). Our previous data show that both Tie2 agonists, H8F and Icos1 upregulated cell migration(9) here, we show that when cells were treated with α5β1mb and H8F, cell migration is significantly reduced to a level similar to PBS control (Fig. 2G-H). Collectively, these data show that α5β1 integrin is critical to Tie2-mediated endothelial cell migration.

### Tie2 activation orchestrates tight junction formation

We previously demonstrated *in vivo* Tie2 activation using a super agonist, Icos1, which accelerated the repair of hemorrhagic vessels after brain injury in mice(9), however, it is unclear how Tie2 regulates this process at the molecular level. Since clustering Tie2 colocalizes with tight junction proteins (Fig. 1E & Fig. S1D), we further evaluated the role of Tie2 in tight junction assembly. To analyze the rate of tight junction recovery under Tie2 signaling, we first disrupted endothelial tight junctions in HUVECs or HBMECs using Latrunculin-A (LatA)(54, 55) before treating them with Tie2 agonist. Briefly, cells were treated with 0.25 μM of LatA for 15 minutes, then LatA was washed off before adding Tie2 antagonist (H3F, Ang2-like) or agonist (H8F, Ang 1-like)(9) for 30 minutes to allow tight junction recovery (Fig. 2I). Our data show that H8F significantly accelerates the recovery levels of both junctional claudin-5 and ZO1 to the cell-cell interphase compared to PBS (30 minutes; >2.5-fold increase), while H3F did not promote any tight junction recovery (Fig. 2I-K). Using live-cell imaging, iPSC-derived endothelial cells (iENDOs) lose junctional ZO1^GFP^ when treated with LatA (Fig. 2L). Once LatA is washed out, cells treated with H8F were able to induce ZO1^GFP^ re-assembly in the cell-cell junctions beginning from 30-90 minutes, while the PBS control has minimal recovery within this time frame (Fig. 2L-M). Tight junction formation plays a crucial role in vascular stability; thus we evaluated the efficacy of H8F for vascular stability in a 72-hour tube assay. Indeed, H8F-treated samples were able to maintain the vascular network significantly longer than the PBS control (Fig. 2N-O). Collectively, our data show that Tie2 activation accelerates tight junction protein assembly to enhance vascular stability.

### Tie2 activation promotes endothelial cell maturation

Endothelial maturation is characterized by the coordinated development of adherens junctions and barrier-forming via endothelial tight junctions(34, 56–60). In figure S3A, we observed that H8F upregulates genes related to tight junction, thus we investigated the role of Tie2 signaling in endothelial cell tight junction formation and maturation during differentiation. We adapted a differentiation protocol to make iPSC-derived endothelial cells (iENDOs) (61) along with Tie2 activation using H8F on day 5 for two days (Fig. 3A). Our data show H8F-treated cultures showed more Tie2 clusters and roughly 15-fold more ZO1-Tie2 cluster formation at the cell-cell junctions by day 8 compared to untreated controls (Fig 3B-C), suggesting that treatment with H8F enhances tight junction formation in developing endothelial cells. We also observed that by day 10, H8F-treated iENDOs are functionally more mature with significant enhancement in cell migration in the wound healing assay compared to control (Fig. 3D-E). H8F-treated iENDOs also performed better in the tube formation assay with higher numbers of nodes, segments, and meshes at the 24-hour time point compared to control (Fig. 3F-G). Together, these results demonstrate that activating Tie2 using H8F accelerates endothelial maturation and functionality, as evidenced by the enriched ZO1 expression at the cell-cell membrane, accelerated cell migration, and enhanced tube formation.

**Figure 3:**
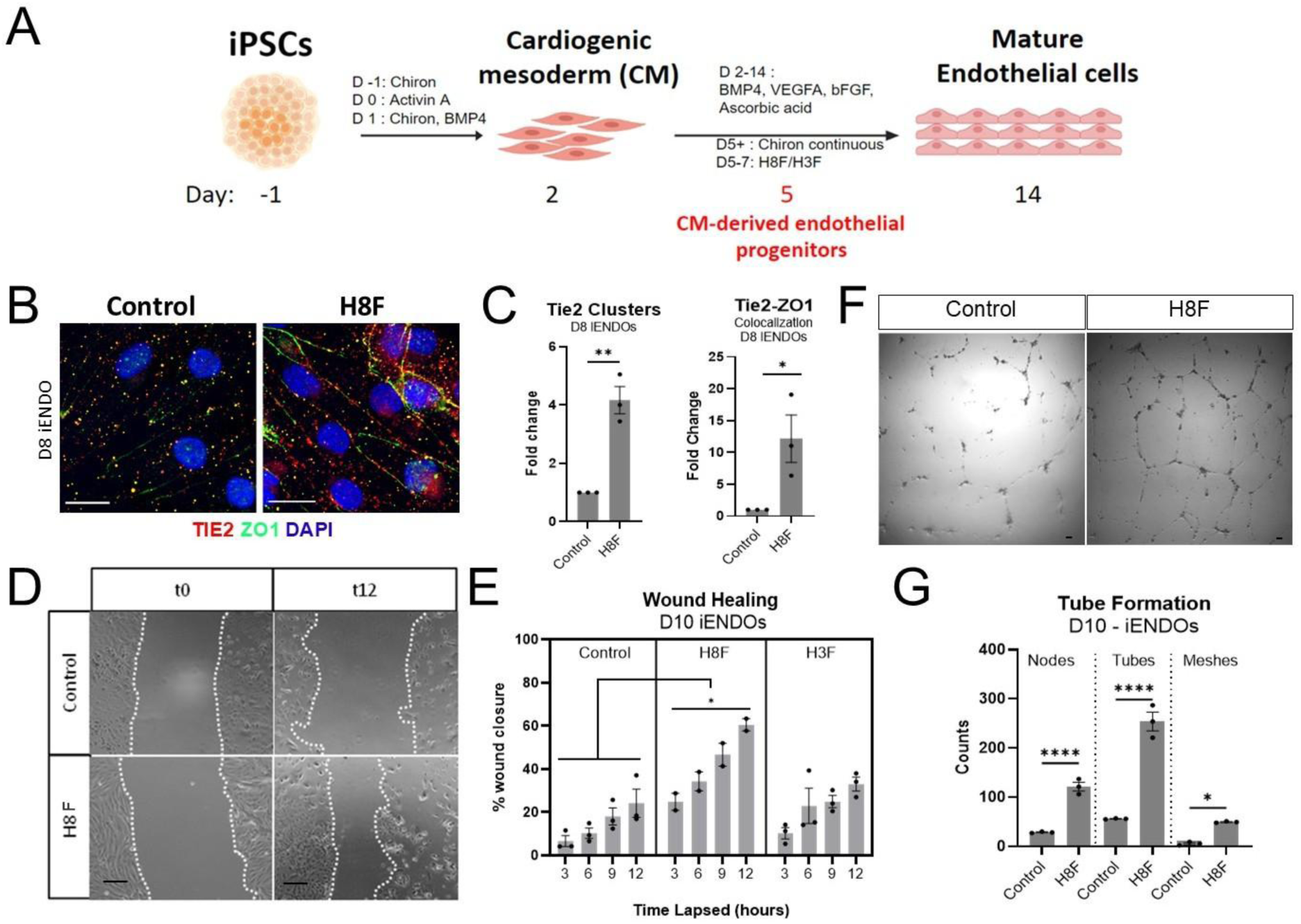
Activating Tie2 promotes tight junction assembly and endothelial cell maturation. **A)** Schematic of endothelial cell differentiation protocol starting from iPSCs, iENDO. **B)** Confocal images of day 8 iENDOs treated with or without 20 nM of H8F on day 5. Cells were stained with ZO1-green, Tie2-red, and DAPI-blue, scale bars are 20 µm. **C)** Quantification of the level of Tie2 clusters and Tie2-ZO1 colocalization found on day 8 iENDOs +/- H8F treatment on day 5 of differentiation. **D)** Brightfield images of day 10 iENDOs in cell scratch assay at 0- and 12-hour timepoints, scale bars are 200 µm. **E)** Quantifications of scratch assay to evaluate wound healing capacity of day 10 iENDOs treated with H8F or H3T at day 5 of differentiation, the least significance level is annotated on the graph for the multiple analysis. **F)** Brightfield images of tube formation assay at the 24-hour timepoint of iENDOs, imaged at 4X magnification, bars are 100 µm. **G)** Quantification of the tubular structures by averaging the number of nodes, meshes, and segments in 24-hour tube formation assay using ImageJ to evaluate the angiogenic capacity of day 14 iENDOs. One-way ANOVA with Bonferroni post hoc test was used to analyze the level of significance, *p*-value < 0.05, 0.01, 0.001, 0.0001 is annotated *, **, ***, and ****, respectively. Each data point is a biological repeat for all experiments.

### Tie2 activation rescues diabetic vasculopathy

Our findings highlight the essential role played by Tie2 in orchestrating tight junction expression and assembly. To determine whether Tie2 activation promotes vascular repair in a disease context, we established an iPSC-derived diabetic vasculopathy (DV) organoid model exhibiting hallmark features of diabetic vascular disease, including increased collagen IV (Col-IV)(62), disrupted endothelial junctions, and nuclear FOXO1 accumulation (Fig. 4A). Mature blood vessel organoids (BVOs) were cultured under non-diabetic conditions, diabetic conditions (high glucose, TNFα, and IL-6), or diabetic conditions followed by 7 days of H8F treatment (Fig. 4A-B). While diabetic conditions disrupted the vascular network, H8F markedly restored vascular architecture (Fig. 4C-D). Three-dimensional analysis showed that diabetes reduced endothelial-pericyte association (VE-cadherin/PDGFRβ colocalization) and increased Col-IV deposition, whereas H8F reversed these changes by decreasing Col-IV (Fig. 4E) and restoring the vascular network (Fig. 4C and S5A).

**Figure 4:**
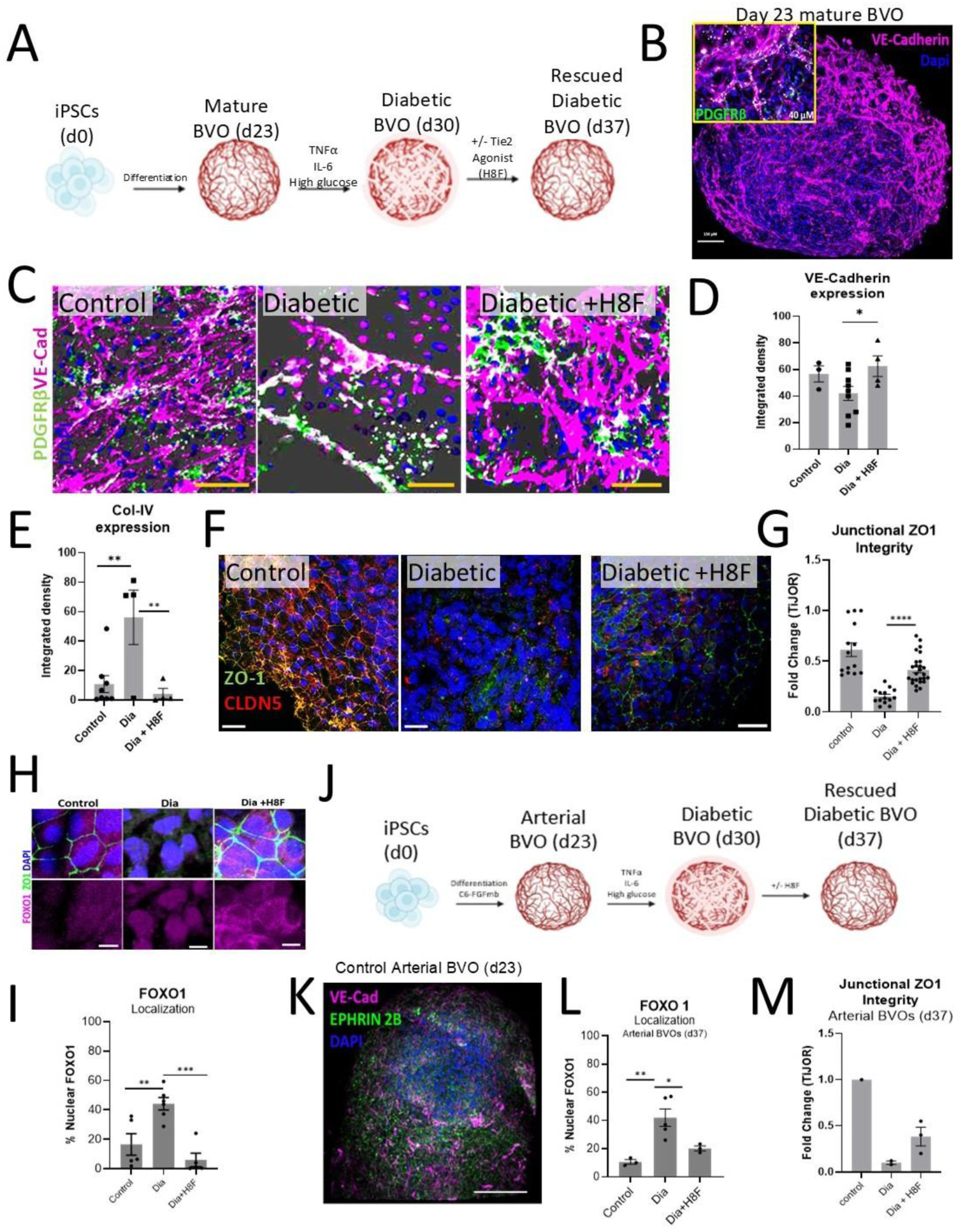
Activating Tie2 rescues diabetic vasculopathy. **A)** Schematic of differentiating iPSCs into 3D blood vessel organoids (BVOs) and treating day 23 mature BVOs with high glucose and cytokines to induce a diabetic condition. On day 30, some BVOs will be treated with H8F at 20 nM. Day 37 BVOs were then fixed and stained with antibodies for confocal imaging. **B)** Day 23 mature BVOs were stained with PDGFRβ, VE-cadherin (CD144), and DAPI, scale bar is 150 µm. **C)** Confocal images of BVOs were processed using Imaris 3D reconstruction of day 37 BVOs under different conditions, scale bars are 50 µm. **D)** VE-cadherin staining in BVOs was quantified for integrated density using ImageJ. Three BVOs were analyzed for all conditions; data points represent sections of the BVO. **E)** Quantification of COL-IV deposition of BVOs at day 37. Immunofluorescence images of BVOs were analyzed using image J to quantify COL-IV integrated intensity, data points represent image sections, three BVO quantified for all conditions. **F)** Representative images of day 37 BVOs were stained with ZO1, Claudin-5, and Dapi for tight junction organization assessment, scale bars are 50 µm. **G)** Quantification of junctional ZO1 integrity using the TiJOR^60^ method. Each data point is an image taken at different z-stacks of the BVOs, at least 2 BVOs were analyzed in each condition. **H)** Confocal images of BVO stained with DAPI and antibodies against FOXO1 and ZO1. **I)** Quantifications of the percentage of cells in BVOs that expressed nuclear FOXO localization, 200-300 cells were counted in each BVO from multiple Z-stacks, each data point is one BVO. **J)** Schematic of iPSC-derived arterial blood vessel organoids using C6-FGFmb during the differentiation step. **K)** Confocal images of day 23 control arterial BVO stained for CD144 (VE-cadherin) and EPHRIN2B to marker arterial blood vessels, scale bars are 150 µm. **L)** Quantifications of nuclear FOXO per cell in arterial BVOs under different conditions. **M)** Quantification of Junctional ZO1 integrity using TiJOR. One-way ANOVA with Bonferroni post hoc test was used to analyze the level of significance, *p*-value < 0.05, 0.01, 0.001, 0.0001 is annotated *, **, ***, and ****, respectively. Each data point is a biological repeat for all experiments (except panels D, E, and G).

Because endothelial barrier disruption is a hallmark of diabetic vasculopathy (20, 63, 64), we next examined tight junction organization. Diabetes caused a profound loss of membrane-localized ZO1 and CLDN5, whereas H8F substantially restored junctional organization, resulting in significantly higher TIJOR scores (Fig. 4F-G). H8F also significantly reduced nuclear FOXO1, a marker of diabetic endothelial dysfunction (65, 66) inflammation(67) and destabilization of blood vessels(37) (Fig. 4H-I).

Since diabetic vasculopathy primarily affects arteries, we generated arterial BVOs using the FGFR1/2 c-splice variant superagonist C6-FGFmb instead of bFGF (68), which increased the arterial marker EphrinB2 (Fig. 4J-K). H8F similarly reduced nuclear FOXO1 in diabetic arterial organoids (Fig. 4L and S5B), demonstrating that Tie2 activation can rescue both general and arterial diabetic vasculopathy. Together, these findings identify H8F as a potent promoter of vascular stabilization and repair under diabetic conditions.

## DISCUSSION

Our study demonstrates that high-valency F-domain ligands cluster Tie2 into distinct signaling complexes that regulate vascular stability through junctional organization and transcriptional control. We identify a Tie2–α5β1 complex containing pCAS and VE-cadherin, linking Tie2 signaling to focal adhesions and adherens junctions that coordinate cell–matrix and cell–cell interactions. This complex is essential for F-domain-induced PI3K/AKT–FOXO1 signaling, supporting endothelial survival and junctional integrity. Consistent with our previous studies, sustained Tie2–α5β1 association prolongs Tie2 residence at the plasma membrane, providing a mechanism for durable vascular stabilization (9, 39).

Previous studies demonstrated that Tie2 promotes adherens junction assembly through VE-cadherin and PECAM (31). Our findings extend this model to tight junctions, showing that Tie2 activation increases the expression and junctional localization of ZO-1, claudin-5, and occludin. Given the role of ZO-1 phase separation in initiating tight junction assembly ((55) and the ability of Tie2 to induce actin remodeling (31), we propose that Tie2 clustering promotes tight junction assembly by coordinating cytoskeletal reorganization with junctional protein recruitment.

Together, these findings support a model in which Tie2 coordinates complementary mechanisms of vascular stabilization (Fig. S4B). At the endothelial–pericyte interface, pericyte-derived Ang1 may engage the Tie2–α5β1 complex to activate AKT, inhibit FOXO1, and support endothelial survival and tight junction gene expression. Through focal adhesion interactions, this complex also promotes endothelial migration at the cell–matrix interface. At cell–cell junctions, Tie2 promotes tight junction protein recruitment independently of α5β1, whereas α5β1 is required for VE-cadherin recruitment. These pathways may cooperate to assemble and stabilize endothelial junctions.

We further demonstrate the therapeutic potential of synthetic Tie2 agonists in a blood vessel organoid model of diabetic vasculopathy. These organoids exhibited collagen IV accumulation, junctional disruption, and nuclear FOXO1 localization.

One week of H8F treatment restored endothelial junctions, reduced collagen IV deposition, normalized FOXO1 localization, and improved vascular organization. These findings show that synthetic Tie2 agonists can reverse established vascular defects in this model, supporting their further investigation as treatments for diabetic vasculopathy and other conditions characterized by endothelial instability.

## AUTHOR CONTRIBUTIONS

HRB, YTZ, and DE conceptualized hypotheses, designed experiments, and prepared the manuscript. YTZ, PL, SY, and CH conducted and analyzed cell assays using F-domain and integrin minibinder conjugated to various scaffolds. XW designed and purified the integrin minibinder. YTZ and XW performed fluorescence polarization assay. YTZ, DE, and AM conducted and analyzed experiments on blood vessel organoids and endothelial differentiation. DE, YTZ, AP, and SS performed sciRNA sequencing analysis. ARH performed the bulk RNA sequencing experiment and YTZ performed the data analysis. DE, YTZ, and AP performed experiments and analysis of arterial blood vessel organoids. HRB, JM, and DB acquired research funding and provided research guidance.

## ACKNOWLEDGEMENTS

This work was supported by grants from the National Institutes of Health DE033016, 1P01GM081619, R01GM097372, and R01GM083867; the NHLBI Progenitor Cell Biology Consortium (U01HL099997; UO1HL099993) SCGE COF220919; DOD PR203328 W81XWH-21-1-0006; and the Stem Cell Gift Funds for H.R-B. NIH/NIDCR #T90DE021984 for YTZ and DE. We thank the Jonathan Cooper Lab at Fred Hutch for sharing pCAS antibodies and guidance in integrin experiments.

## DATA AVAILABILITY STATEMENT

The bulk RNA sequencing data presented in the study is deposited in the GEO Database repository and accession number will be provided upon publication.

## COMPLETING INTERESTS

An International PCT Patent Application, entitled “De novo designed Tie2 mini binder acting as an agonist or antagonist for Tie2 receptor,” was filed, serial number PCT/US2025/058945. This patent encompasses the matter of use claim for the therapeutic application of H8F in diabetic vasculopathy.

## MATERIALS AND METHODS

### Cell culture

Human umbilical vein endothelial cells (HUVECs) were purchased from Lonza (Catalog #: C2519A) and cultured in EGM2 media described previously^9^. Briefly EGM2 consist of 20% fetal bovine serum (BioWest, S1620), 1% penicillin–streptomycin (Gibco, 15140122), 1% GlutaMAX (Gibco, 35050061), 1% ECGS (endothelial cell growth factor), 1 mM sodium pyruvate (Gibco, 11360070), 7.5 mM HEPES (Gibco, 15630130), 0.08 mg/ml heparin (Fisher BioReagents, 904108-1), 0.01% amphotericin B (Gibco, 15290018), a mixture of 1x RPMI 1640 (Gibco, 1875093) with and without glucose (Gibco, 11879020) to reach 5.6 mM glucose concentration in the final volume. Media was filtered through a 0.2-μm filter. HUVECs were expanded and serially passaged to reach passage 4 before cryopreservation. Human brain microvessel endothelial cells (HBMECs) were purchased from Cell Systems (catalog # ACBRI 376) and cultured in EGM^TM^-2 MV Microvascular Endothelial Cell Growth Medium-2 BulletKit^TM^ (catalog # CC-3202, Lonza). Both cell lines were cultured until passage 7 or 8 before 16 hours of serum starvation in low glucose DMEM (catalog # 11885084, Gibco) before experiments.

hiPSCs WTC-11 human (Coriell Institute, #GM25256) and TJP1-GFP WTC11 hiPSCs line are generated by the Allen Institute for Cell Catalog Science (AICS-0023 cl.20), purchased from Coriell Institute, expanded in the Ellison Stem Cell Core at the University of Washington (UW-AICS-0023 cl.20) - were cultured with mTeSR™1 media (STEMCELL, #85850) on plates coated in 2.5% Matrigel (Corning, #356231) until differentiation experiments.

### Western blot for signaling analysis

Serum-starved HUVECs were treated with synthetic ligands for 15 minutes at 37 degrees Celsius. Treatment was then removed, and cells were washed once with 1x PBS before adding the lysis buffer described previously^9^. Lysates were then collected in tubes, and 4X Laemmli sample buffer (BioRad, #161-0747) was added before boiling at 95 degrees Celsius for 10 minutes. 30 µL of protein sample per well was loaded and separated on a 4–10% SDS–PAGE gel for 30 min at 250 Volt. The proteins were then transferred onto a nitrocellulose membrane for 12 min using the semi-dry turbo transfer Western blot apparatus (Bio-Rad, USA). Post-transfer, the membrane was blocked in 5% bovine serum albumin for 1 hour. After blocking, the membrane was probed with the respective primary antibodies and incubated at 4°C, overnight on a rocker: pAkt-S473 (Cell Signaling, 9271S) at 1:2,000 dilution; pERK1/2 p44/42 (Cell signaling, 4370S) at 1:1,000 dilution; Tie2 (Cell Signaling,4224S) at 1:1,000 dilution; α5 integrin (Millipore, AB1928) at 1:1,000; and β-Actin (Cell Signaling, 3700S), S6 (Cell Signaling, 2217S). The next day, the membranes were washed with 1X TBST (3 times, 10 min intervals) before adding the respective HRP-conjugated secondary antibody (Bio-Rad, USA) at a 1:10,000 dilution and incubated at room temperature for 1 hour. Following washes for pAKT(S473), the membrane was blocked in 5% milk at room temperature for 1 hour and then incubated with its respective HRP-conjugated secondary antibody (1:2,000) prepared in 5% milk for 1 hour. After secondary, the membranes were washed with 1× TBST (3 times, 10 min interval), developed using Chemiluminescence developer, and imaged using Bio-Rad ChemiDoc Imager. Quantifications were done by calculating the peak area for each band on Western blot images using ImageJ. The peak area of the target proteins was divided by the peak area of the housekeeping proteins. All signaling levels are normalized to H8F levels as an internal positive control.

### Protein conjugations

Design protein component genetically fused with SpyCatcher was combined with binder genetically fused with SpyTag (F-domain-st, Tmb-st, α5β1mb, or GFP-st) at a 1:1 – 1.5 molar ratio for 4 hours at room temperature or overnight at 4°C on nutation. SDS-electrophoresis gels were used to evaluate conjugation efficiency. All experiments were done using constructs that have reached at least 90% conjugation efficiency. Two-dimensional sheet assembly was done according to the previous protocol^26^. Briefly, A-GFP, A-(Fd or mb), and B domains were combined at a 1:4:5 molar ratio, respectively, and incubated at room temperature for 2 hours or overnight at 4 degrees C on nutation before experiments. Protein constructs were run on SDS gels and stained with Coomassie blue to confirm conjugation and assembly Figure S6.

### Immunofluorescence analysis of Tie2 clusters

Serum-starved HUVECs or HBMECs were treated with Fd-sheets or α5β1mb-sheets at 50nM of binding domain for 10-30 minutes in low glucose DMEM before fixation with 4% PFA. The fixed cells were washed three times at 5 minutes each before blocking for 1 hour with 3% BSA (VWR, 0332-500G) and 0.1% Triton X-100 (Sigma, T9284-500ML) in PBS while on nutation. The cells were then incubated with primary antibody diluted at 1:100 in blocking agent overnight: Tie2 (Cell Signaling, catalog #4224S) and α5 integrin (Millipore, 1928), active β1 integrin 9EG7 (BD Biosciences, 550531), inactive β1 integrin mAb13 (BD Biosciences, 552828), phospho-CAS, CAS (Cell Signaling, 4011S), CD144/VE-cadherin (Cell Signaling, 2500S), ZO1 (Invitrogen, 33-9100 or 61-7300), claudin-5 (abcam, ab15106), and occludin (Invitrogen, 710192). After the primary antibody, the cells were washed three times for 5 minutes each with 1X PBS while on nutation. The cells were then incubated with secondary antibodies at 1:200 (1:100 for Tie2) and phalloidin at 1:100 (Invitrogen, A12380) diluted in blocking agent for 1 hour and 20 minutes at 37° C. Secondary antibodies were then removed, and cells were washed three times at 10 minutes each with PBS on nutation. Coverslips were sealed using VECTASHIELD plus DAPI (Vector Laboratories, H-2000-2) upside-down on glass slides for analysis in confocal (Leica SP6 or SP8) or super-resolution (Delta vision OMX SR) microscopy.

### Fluorescence polarization assay

Fluorescence polarization assays were performed in low-binding black 96-well microplates using an optimized physiological buffer containing 20mM tris, 150mM NaCl, 1mM Ca2+, and 1mM Mg2+, 0.01% Tween-20. Reaction mixtures containing the fluorescently labeled tracer FIT-C tagged RGD peptide and a5b1 integrin ectodomain protein were mixed before adding H8F to the mixture. Mixtures were incubated in the dark at room temperature for 20 minutes. Fluorescence emissions were measured using polarized excitation, and binding affinity or inhibition was quantified in millipolarization (mP) units based on parallel and perpendicular emission intensities.

### FOXO1 nuclear exclusion assay

The FOXO1 localization analysis was performed according to the protocol described in Zhao et al 2021. Briefly, serum-starved HUVECs were treated with Tie2 binders for 30 minutes in low-glucose DMEM before fixation with 4% PFA. The fixed cells were washed three times at 5 minutes each before blocking for 1 hour with 3% BSA (VWR, 0332-500G) and 0.1% Triton X-100 (Sigma, T9284-500ML) in PBS. After blocking, the primary antibody FOXO1 (Cell Signaling, 2880) was diluted in the blocking agent overnight at a 1:100 ratio. After the primary antibody, cells were washed three times at 5 minutes each with 1X PBS. The cells were then incubated with secondary antibodies at 1:200 dilution in blocking agent for 1 hour and 20 minutes at 37° C. Then, cells were washed three times at 10 minutes each with PBS. Coverslips were sealed using VECTASHIELD plus DAPI (Vector Laboratories, H-2000-2) upside-down on glass slides for analysis in confocal using microscopy Leica SP6 or SP8.

### SiRNA knockdown of *TEK* or *ITGA5*

Passage 7 HUVECs (100,000 cells/1 mL) were first transfected with 40 nM siRNA using Lipofectamine Transfection Kit (Invitrogen, 13778-075) in 1 mL EGM2 medium suspension without antibiotics. Cell suspensions were then seeded on 0.1% gelatin-coated 35-mm plate (for sheet experiments, cells were seeded onto coverslips coated with gelatin). After 18 hours of siRNA incubation, media were changed with fresh EGM2 to allow cell recovery for 6 hours before 16-hour serum starvation with low glucose DMEM. Starved HUVECs were treated with scaffolds at 20 nM of F-domain concentrations for 15 minutes before protein collection for western blot analysis or 30 minutes for sheet experiments before PFA fixation. siRNAs targeting *TEK* (siRNA: M-003178-03-0005) or *ITGA5* (siRNA: L-00800300-0005) mRNA were purchased from Dharmacon™.

### Cell migration assay

Scratch assay was performed according to previous protocols with modifications^9,39^. Before seeding cells, one line is drawn on the bottom of the plate perpendicular to the scratch to track imaging spots. HUVECs or day 10 iENDOs were seeded on 0.1% gelatin-coated surface and cultured in homemade EGM2 media (HUVECs) or Lonza EGM2 Bullet Kit (iENDOs) until confluent. Once cells reach 80-90% confluency, a scratch is made on the center of the plate, and then washed once with PBS, and incubated in low-glucose DMEM supplemented with 2% FBS +/- treatment for 12 hours. HUVECs were imaged at 0 and 12-hour time points. For iENDOs, the wound area was imaged at 0-, 3-, 6-, 9-, and 12-hour time points. Images were taken above and below the drawn line to get two images per well. ImageJ was used to quantify the images at all timepoints and in pixel scale. The percent change of wound area was calculated as below:

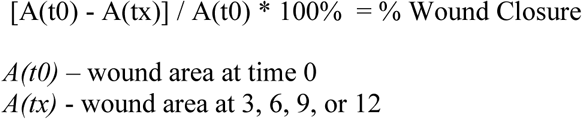

### Bulk RNA sequencing

HUVECs were grown till 80% confluence and then serum starved for 6 hours in DMEM LG media before administrating 10nM of H8F with or without 100nM of α5β1mb. Cell lysates were collected using DNA/RNA Shield lysis buffer (Zymo, R1100-50) and sent out for bulk RNA sequencing at Plasmidsaurus. FASTq files from sequencing results were uploaded onto Partek processing and analysis. DESeq2 were performed on the processed data to compare differentially expressed genes between H8F + α5β1mb vs H8F. List of downregulated genes in H8F + α5β1mb samples were analyzed for expression level using heatmap and pathway enrichment.

### Tight junction disruption using Latrunculin-A

To evaluate the effect of Tie2 activation on tight junction re-assembly, endothelial cells were treated with an actin inhibitor, Latrunculin-A (Cayman, 10010630), to disturb tight junction proteins^41^ before F-domain scaffold administration. Briefly, confluent HUVECs or HBMECs seeded on coverslips were treated with 0.25µM of LatA diluted in growth media for 30 minutes. After 30 minutes, LatA was washed out with PBS before adding F-domain scaffolds at 100 or 200 nM diluted in LG DMEM + 10% FBS for 30 minutes or 2 hours. Post-treatment, cells were fixed with 4% PFA for immunofluorescence staining with antibodies (detailed protocol in the immunofluorescence section) to visualize ZO1 (Invitrogen, 1:100) or Claudin-5 (abcam, 1:100). Phalloidin (1:100, Invitrogen, A12380) was added during secondary staining. Stained cells were mounted on slides using VECTASHIELD plus DAPI. Phalloidin stains were used to find cell colonies with good contact before switching to ZO1/CLDN5 stains to count junctional tight junction assembly. At least 100 cells were counted per coverslip.

### Tube formation assay

Tube formation assay was done using modified protocols^9,62^. For iENDO tube formation assay, day 10 iENDOs were lifted with Accutase and replated at 150,000 cells/well in 24-well tissue culture plates coated with 100% Matrigel. 150 μL of Matrigel was solidified for 30 minutes at room temperature before cell seeding. Cells were suspended in 350 μL of Low Glucose (1g/L) DMEM + 0.5% FBS. The cells were incubated for 24 hours to allow tube formation before imaging using a brightfield microscope at 10X magnification; 10 images were taken per well at random spots and then quantified by counting the number of meshes, tubes, and nodes. H8F samples were pre-treated as described above on day 5 of differentiation.

For vascular stability analysis, HUVECs were seeded onto 100% Matrigel with or without 100 nM of H8F for 24 hours to allow vasculogenesis. At the 24-hour time point, fresh media were added to remove H8F and continue to incubate up to 72 hours. At the 72-hour timepoint, tubule networks were imaged and quantified using the ImageJ angiogenesis analyzer plugin. Vascular stability was counted by averaging the number of meshes, tubes, and nodes. Vascular stability was normalized to the PBS control.

### iPSC-derived endothelial cell differentiation protocol

iPSC-derived endothelial (iENDO) cells were differentiated as previously described(61). Briefly, human induced pluripotent stem cells WTC-11 or WTC-11 TJP1-GFP cells were seeded at 2.5 x 10^5^ cells/well on 24-well plates coated with 2.5% growth factor-reduced Matrigel [Corning, #356231] and cultured in mTeSR1 stem cell medium [StemCell Technologies, #85850] supplemented with 1µM GSK3-Inhibitor (CHIR99021) [Cayman Chemicals, #13122] for 24 hours. On the first day of differentiation (D0), the media was replaced with cardiogenic mesoderm media consisting of RPMI 1640 Medium [Thermo, #11875093] supplemented with 1X B27 without insulin [Fisher Scientific, #A1895601], 100 ng/mL Activin A [PeproTech, #120-14P], and 2.5% Matrigel for 17 hrs. The next day, the media was replaced with RPMI supplemented with 1µM of GSK3-Inhibitor (CHIR99021), B27 without insulin, and 5 ng/mL bone morphogenetic protein-4 (BMP-4) [R&D systems, #314-BP-010] for 24 hours. On Day 2 of differentiation, cells were washed with 1X PBS and media was replaced with vascular differentiation media consisting of StemPro [Thermo Fisher, #10639011] supplemented with 1X GlutaMAX, 1X Penicillin-Streptomycin, 300 ng/mL vascular endothelial growth factor (VEGFA165) [R&D systems, #293-VE-050], 10 ng/mL BMP-4, 5 ng/mL FGF2 (10nM of C6 were used instead to make arterial blood vessel organoids), 50 ug/mL Ascorbic Acid [Sigma-Aldrich, #A8960], and 40 µM monothioglycerol (MTG) [Sigma-Aldrich, #M6145]. On Day 5, cells were dissociated with Accutase [Thermo, #A1110501] and replated 6,667 cells/cm^2^ in tissue culture wells (assay-specific sizes) coated with 0.1% gelatin in EGM2 Bullet Kit endothelial media [Lonza, #CC-3121] supplemented with 20 ng/mL VEGF, 20 ng/mL FGF2, and 1µM GSK3-Inhibitor (CHIR99021). Media was replaced every 48 hours until the final harvest on Day 14. For samples that were treated with H8F, this was added at 20nM during the replating step at d5 and left for 48 hours.

### Diabetic blood vessel organoids

This protocol was adapted from a previous version^44^. To create iPSC-derived vascular organoids, WTC-11 hiPSCs were passaged, counted, and resuspended in differentiation media (DMEM/F12, 20% KOSR (Gibco, #10828028), 1X GlutaMAX, 1X NEAA (Gibco, #11140050) supplemented with 50μM Y-27632 [Tocris Bioscience, #1254] and distributed into Ultra Low Attachment 6-well culture plates (Corning, #07-200-601) at a density of 3 x 10^5^ cells/well to allow spheroid formation. At day 3, cells were fed with fresh differentiation media supplemented with 12μM CHIR99021. At days 5, 7, and 9, cells were given fresh differentiation media supplemented with 30ng/ml recombinant BMP-4, 30ng/ml recombinant VEGFA165 (R&D Systems, #293-VE), and 30ng/ml recombinant FGF2. For Days 11-15, the glucose in the differentiation media was reduced to 2 g/L glucose from 3.15 g/L by using SILAC Advanced DMEM/F12 (Gibco, #A2494301) supplemented with 147.5 mg/mL L-Arginine, 91.25 mg/mL L-Lysine, 2 g/L D-Glucose, 20% KOSR, 1X GlutaMAX, and 1X NEAA. On Day 11, cells were given fresh differentiation media supplemented with 30ng/mL VEGFA165, 30ng/mL FGF2, and 10 μM SB431542 (Miltenyi, #130-106-275). On Day 13, cells were transferred to Ultra Low Attachment U-Bottom 96-well plates (Corning #CLS3474-24EA) at 1 organoid per well, embedded in 1:1 Matrigel: Collagen I matrix, and overlaid with differentiation media supplemented with 100 ng/mL VEGFA165 and 100 ng/mL FGF2. From Days 15 to 23, the glucose in the differentiation media was reduced to 1g/L glucose (physiological glucose) by making it as described for Days 11-15, but supplemented with 1 g/L glucose. Cells were fed on Day 15 and every 3 days thereafter until Day 37.

To create diabetic vascular organoids, at d23 (when organoids are mature), organoids were fed with media described above, but containing 4.5g/L D-glucose and 1nM each of recombinant TNFα and IL-6 inflammatory cytokines. The media was changed every 3 days until harvest at d37. For organoids treated with H8F, organoids were first cultured in diabetic media for a period of 7 days (through d30) and then treated with 20nM H8F through d37.

### Immunofluorescence staining of diabetic blood vessel organoids

All staining steps were done with a rocker to allow adequate mixing. Harvested BVOs were fixed in 1X PBS with 4% PFA for 15 minutes. Fixed BVOs were washed 3 times with PBS at 5 minutes each and then blocked in 3% BSA + 0.1% triton for 2 hours before adding primary antibodies (Claudin5 AbCam#15106, VE-cadherin CST#2500, PDGFRβ CST#3169, Col-IV Millipore#AB769, FOXO1 CST#2880, ZO1 Invitrogen#33-9100) at 1:100 dilution and incubated overnight in 4°C. After removing the primary antibodies, the BVOs were then washed 3 times with PBS for 5 minutes each. Secondary antibodies and DAPI were diluted in blocking solution at a 1:200 dilution and incubated overnight at 4 degrees C. After removing secondary antibodies, BVOs were washed 3 times with PBS for 5 minutes each. BVOs were then transferred to a glass slide lined with rubber holders to hold one BVO per well and embedded with 20uL of Vector Shield. BVOs were then imaged using the Leica SP8 confocal microscope.

For Col-IV and VE-Cadherin immunofluorescence staining quantification, about 50 slices in the z-stacks of BVO were collapsed into one image using ImageJ max projection. Integrated density was used to analyze fluorescence intensity of the images.

### Tight junction scoring method - TiJOR

Quantification of junctional ZO1 integrity in the BVOs using the TiJOR^60^ method. Briefly, about 50 slices in the z-stacks of BVOs into one image. Multiple stacked images of one BVOs were analyzed using the TiJOR method to analyze junctional ZO1 organization. Raw scores were then normalized to control and plotted as fold change.

## SUPPLEMENTAL FIGURE LEGENDS

**Figure S1:**
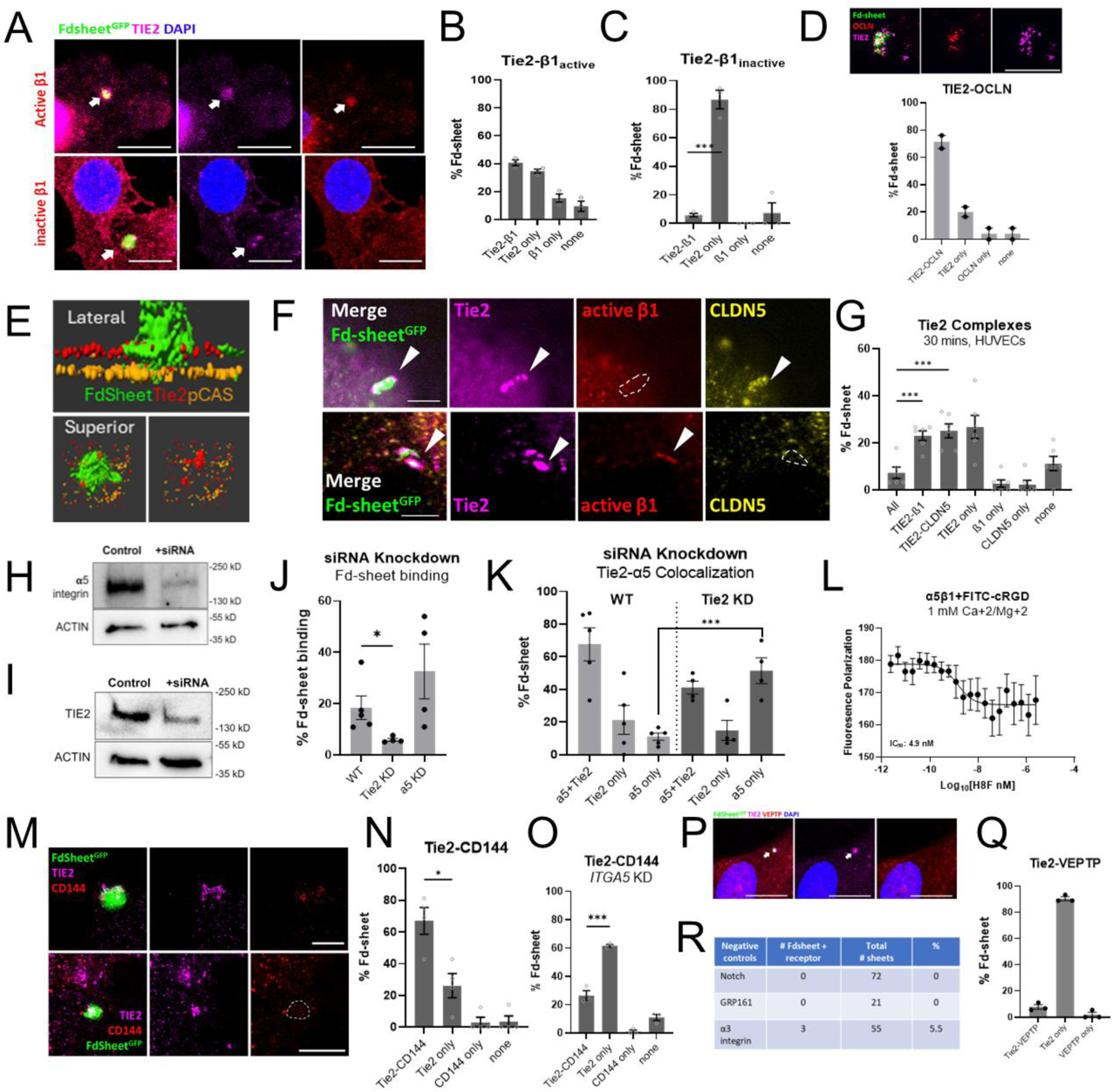
**A)** Representative confocal images of Fd-sheet cluster Tie2 receptors that recruit and activate integrin shown by the presence of active β1 subunit not inactive β1 staining, scale bars are 10 µm. **B-C)** Quantification of Tie2-activeβ1 or Tie2-inactiveβ1 colocalization under the Fd-sheet. **D)** Top: representative OMX images and quantification of Tie2-occludin localization under the Fd-sheet, scale bar is 5 µm. Bottom: quantification of Tie2-occludin colocalization under the Fd-sheet. **E)** Zoom out Imaris images of Tie2-pCAS colocalization under the Fd-sheet in Figure 1C. **F)** Representative confocal images of Tie2 colocalizing with either activeβ1 or claudin-5 under the Fd-sheet using 5 color immunofluorescence imaging, scale bars are 5 µm. **G)** Quantification of panel J summarizing the different Tie2 complexes found under the Fd-sheet. **H-I)** Western blot staining of HUVECs pre-treated with 40nM of siRNA against ITGA5 or TEK to knockdown (KD) α5 integrin or Tie2 expression, respectively, before starvation for Fd-sheet experiments. **J)** Quantification showing the reduction of Fd-sheet binding to Tie2 KD compared to wildtype, α5 KD did not affect Fd-sheet binding to HUVECs. **K)** Quantifications showing the reduction of Tie2-α5 cluster and the significant increase in α5-only cluster under the Fd-sheet in Tie2 KD compared to wild type. **L)** Quantification of fluorescence polarization assay showing Fd binding to integrin. **M)** OMX images of Tie2-CD144(VE-cadherin) colocalization in wild-type (top) vs α5 KD (bottom) cells, scale bars are 5 µm. **N)** Quantifications showing the percentage of Tie2-CD144 colocalization under the Fd-sheet in wildtype and **O)** the significant reduction of Tie2-CD144 colocalization under the Fd-sheet in α5 KD cells. **P-Q)** Clustering Tie2 with Fd-sheet does not recruit VE-PTP, scale bars are 10 µm. **R)** Summary showing that clustering Tie2 does not recruit unrelated surface receptors, Notch, GPR161, and α3 integrin.

**Figure S2.**
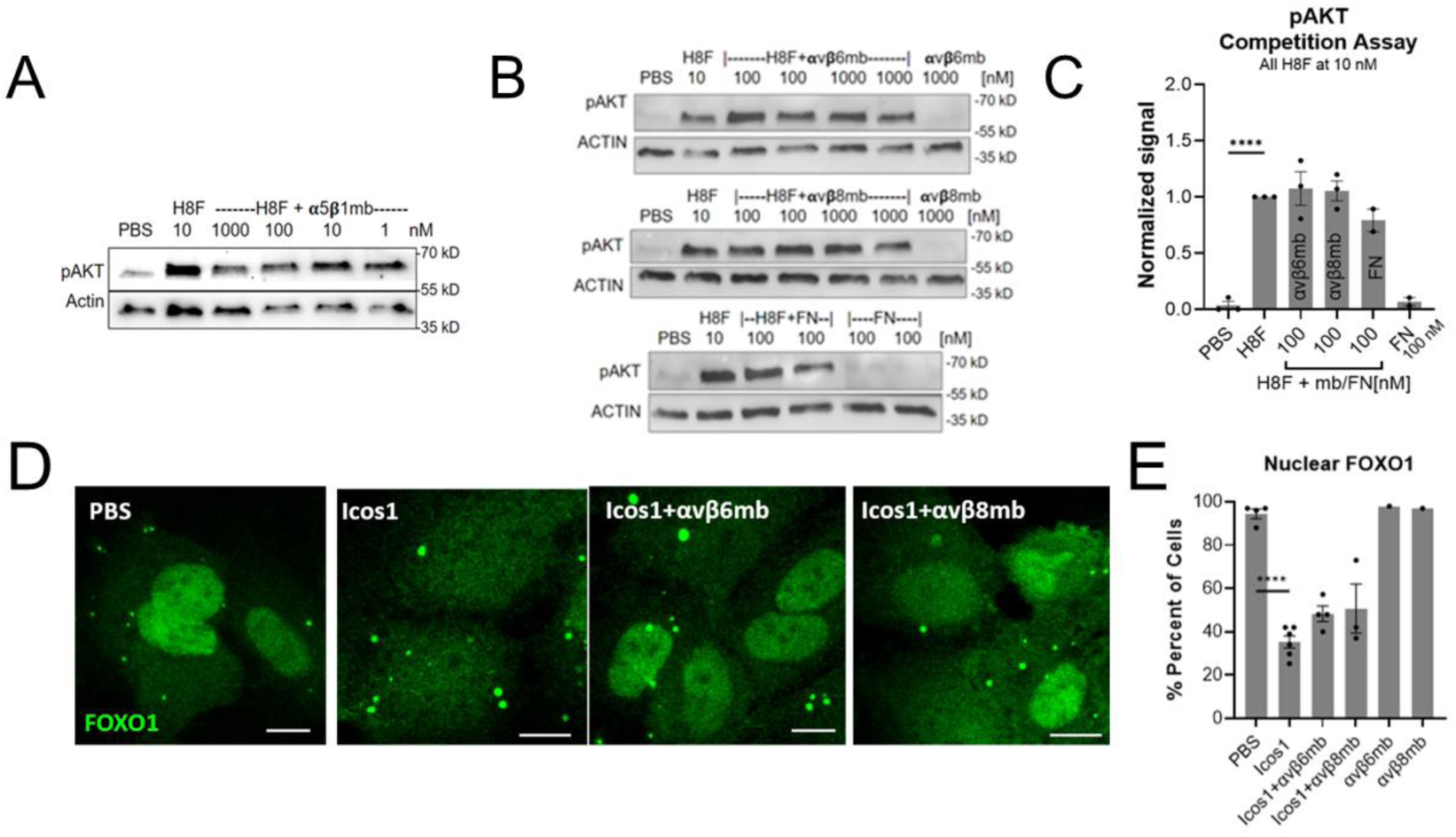
**A)** Western blot images of pAKT inhibition upon treatment with H8F and α5β1mb in HUVECs. **B)** Western blot images of pAKT activation of HUVECs treated with 10 nM of H8F +/- αvβ6mb, αvβ8mb, or fibronectin (FN) at 100-1000 nM. **C)** Quantifications of pAKT/S6 level normalized to 10 nM of H8F in competition experiments. **D)** Immunofluorescence images of FOXO1 staining upon α5β1mb administration, scale bar is 5 µm. **E)** Quantification of nuclear FOXO1 in HUVECs upon the administration with Icos1 +/- αvβ6mb or αvβ8mb, scale bars are 5 µm.

**Figure S3.**
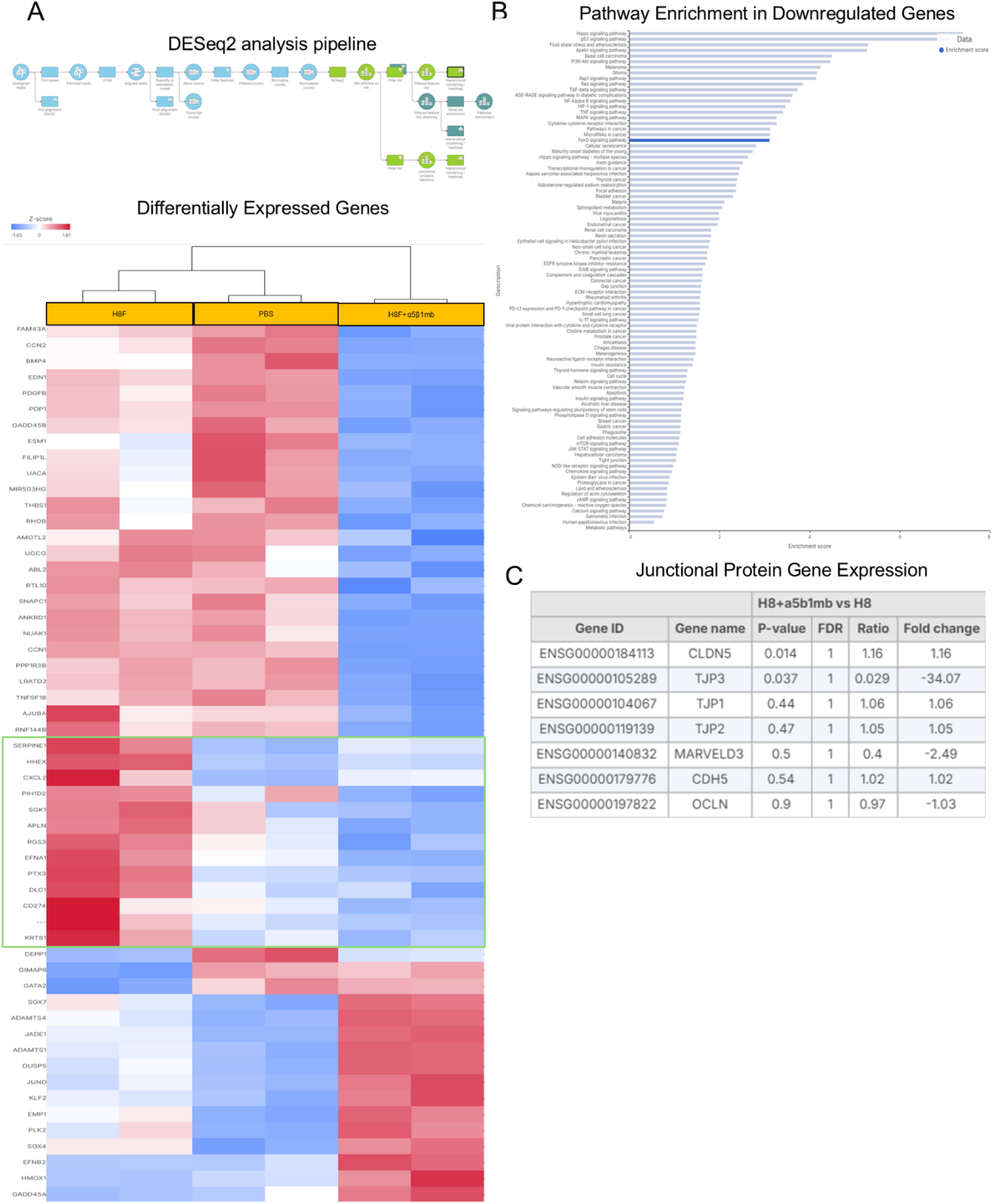
**A)** Top - bulk RNA sequencing data were processed and analyzed in Partek using DESeq2 for differentially expressed genes. Bottom - heatmap of significantly differentially expressed genes comparing H8F + α5β1mb to H8F. The green box are genes showing downregulated genes in both H8F + α5β1mb and PBS compared to H8F, also shown in figure 2E. **B)** Significantly downregulated genes in H8F + α5β1mb were analyzed for pathway enrichment. **C)** Level of gene expression of junctional proteins of interest.

**Figure S4.**
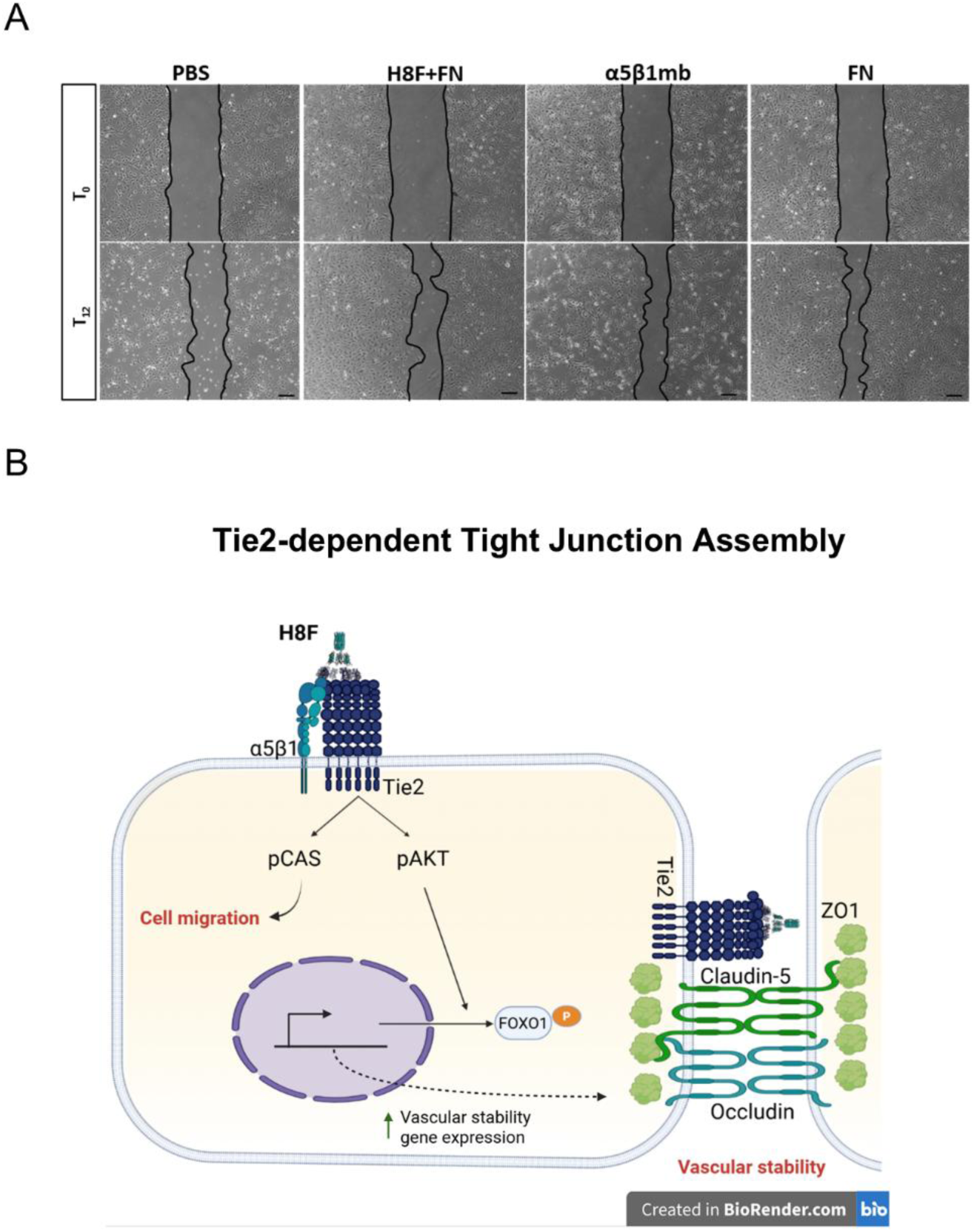
**A**) Brightfield images of HUVECs at 0 and 12 timepoints in scratch assay with 20 nM of H8F +/- FN 100nM, 20 nM of α5β1mb or PSB control, scale bars are 200 µm. **B)** A schematic of the proposed multimodal functions of Tie2 in vascular stability. Tie2 forming two classes of Tie2 complexes and their respective cellular functions, created in RioRender.com.

**Figure S5.**
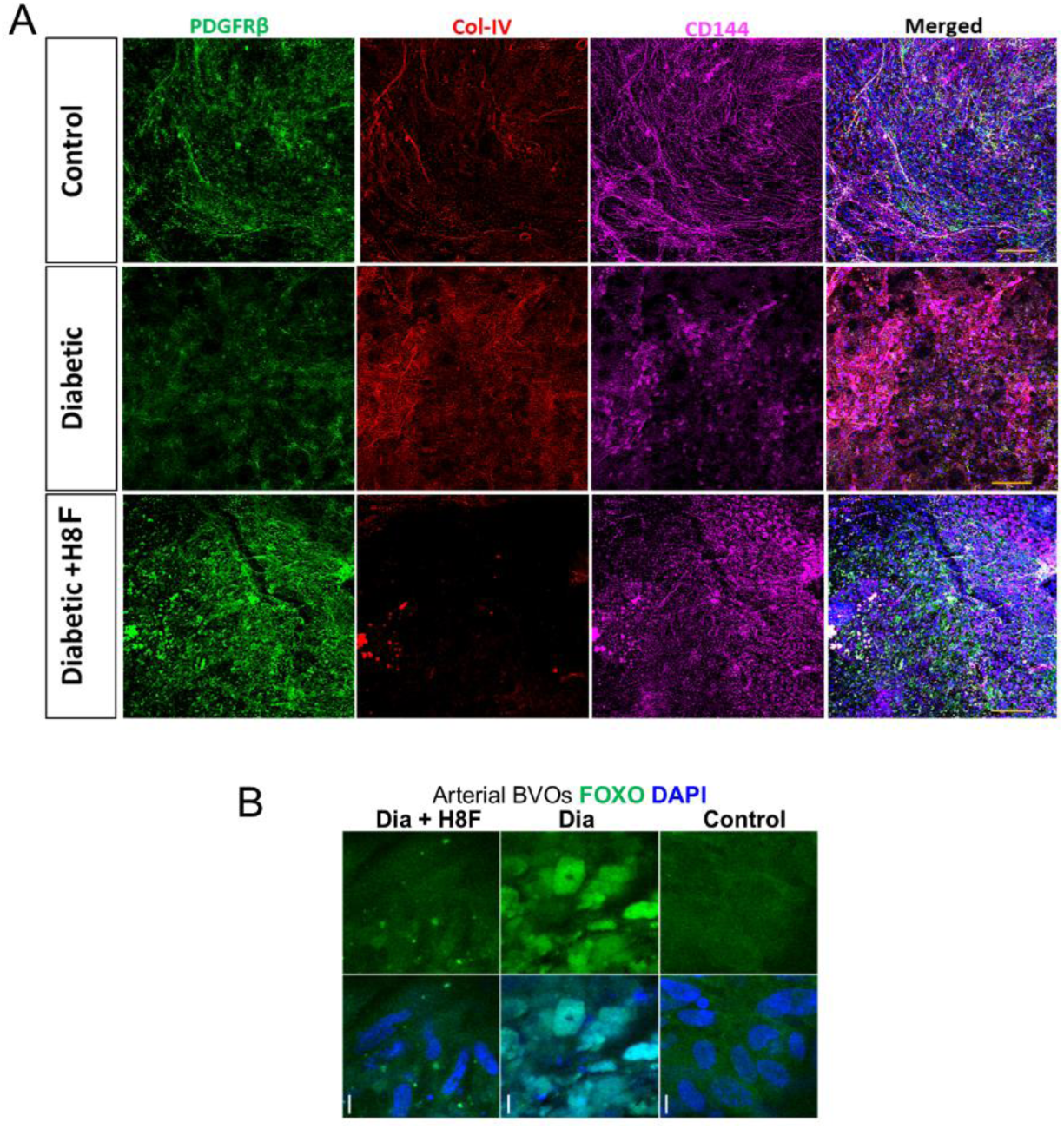
**A)** Confocal images of day 32 BVOs at different conditions stained with PDGFRβ, Col-IV, CD144/VE-cadherin, and Dapi. Scale bars are 150 µm. **B)** Confocal images of FOXO1 stains in day 37 arterial BVOs under different conditions, scale bars are 5 µm.

**Figure S6.**
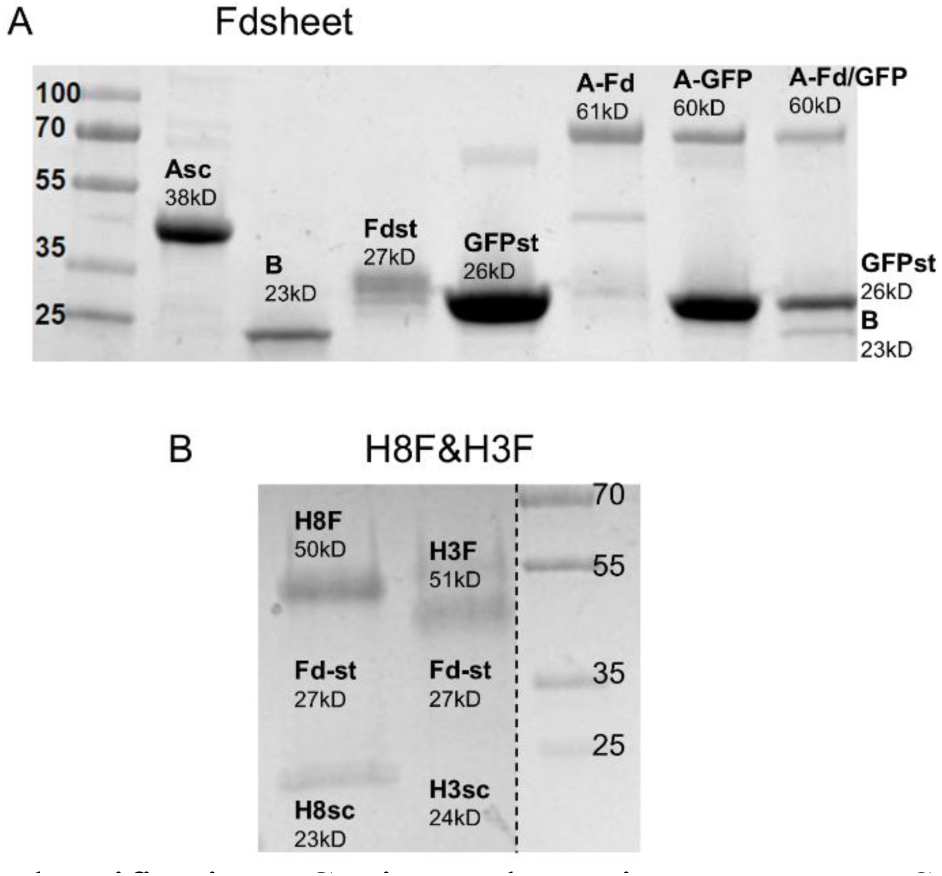
Conjugation efficiency and verifications. Conjugated proteins were run on SDS gels and stained with Coomassie Blue to visualize protein subunits in each scaffold. **A)** F-domain-SpyTag (Fdst) conjugated to the A domain-SpyCatcher (Asc) to make A-Fd and then assembled with A-GFP and B domain to make Fd-sheet. **B)** Fdst were conjugated to H8-spycatcher (H8sc) or H3-spycatcher (H3sc) to make H8F or H3F.

